# The dynamic and structural properties of axonemal tubulins support the high length stability of cilia

**DOI:** 10.1101/451351

**Authors:** Ron Orbach, Jonathon Howard

## Abstract

Cilia and flagella, which play essential roles in cell motility, sensing and development, contain at their core a microtubule-based structure called axoneme. The axoneme, which contains nine doublet and two central microtubules, is highly stable and its length is precisely-controlled. We have asked whether the axonemal tubulins contribute to this length stability. Towards this end, we used a novel procedure to differentially extract the tubulins from the different axoneme components, and characterized their dynamic properties and post-translational modifications (PTMs). We found that their dynamic properties are consistent with the greater stability of axonemes. Despite the differences in PTMs between the axonemal components, we found no significant differences in their dynamic properties. Unexpectedly, the reconstituted ciliary microtubules exhibit curved protofilaments at their growing tip, that may correspond to the fluctuating GTP-cap hypothesized to exist. Thus, our study provides new insights into the growth, stability and the role of PTMs of ciliary tubulin.

#### Abbreviations

MTs: microtubules;
PTMs: post-translational modifications;
PFs: protofilaments;
CP: central pair;
MAPs: microtubule-associated proteins;
IRM: interference reflection microscopy;
TEM: transmission electron microscopy;
cryo-EM: cryo-electron microscopy;
MSD: mean squared displacement;
GMP-CPP: guanosine-5’-[(α,β)methyleno]triphosphate

---

Eukaryotic cilia and flagella, terms often used interchangeably, are evolutionarily conserved hair-like organelles^1^ that play key roles in cell motility, mechanical and chemical sensing, and development^2,3^. The core structure of the cilium, the axoneme, has a “9+2” microtubule (MT) architecture: two singlet MTs, known as the central pair (CP), and nine surrounding doublet MTs^4,5^. The singlet MTs are hollow cylindrical structures formed by 13 protofilaments (PFs) of the ab-tubulin heterodimers. The doublet MTs consist of the A-tubule, which forms a complete MT with 13 PFs, and the B-tubule, which is attached to the A-tubule, and forms an incomplete MT structure with 10 PFs^6^. Interestingly, the MTs in the axoneme are very stable against depolymerization compared to cytoplasmic MTs. Behnke & Forer identified four classes of MTs based on their resistance to high temperature: cytoplasmic MTs < CP < B-tubule < A-tubule^7^. This result was further supported by another study using a surfactant to depolymerize doublet MTs^8^.

MTs are dynamic polymers whose ends switch stochastically between phases of slow growth and rapid shrinkage, in a process called dynamic instability. The switch from growth to shrinkage is called catastrophe and the reverse switch from shrinkage to growth is called rescue^9^. This dynamic nature of MTs is a key element that drives various cellular processes including cell architecture, division, and motility^10-12^. In cilia, however, dynamic instability is suppressed, and instead, assembly and disassembly are in a steady state, through mechanisms that are not understood, leading to remarkable length stability^13^.

In this study, we have asked whether the tubulins from the various axoneme components have different dynamic and functional properties that contribute to the length stability of axonemes. One of the important questions we address is the role of post-translational modifications (PTMs), which have been proposed to regulate many properties of cilia, such as length^14^, stability^15-18^, and motility^19-21^, but have not been studied in the context of MT dynamics. This question remained open until now due to the difficulty in purifying functional tubulins from the axoneme.

## Results

### Purification of axonemal tubulin

We decided to use the unicellular biflagellate green alga *Chlamydomonas reinhardtii* to study flagellar MTs in part because any molecular heterogeneity of their MTs is due only to PTMs. *C. reinhardtii* has two genes that encode a-tubulin and two genes that encode b-tubulin^22^. Each pair of genes specifies an identical protein product and therefore only a single type of tubulin heterodimer is synthesized^23,24^. To overcome the challenge of extracting functional tubulins, we decided to use the Hofmeister series^25^, which classifies the effect of different ions on the solubility and stability of proteins^26^. We searched for a salt that destabilizes the axoneme structure, but does not cause tubulin denaturation and the loss of its ability to repolymerize. Purified axonemes were incubated with different salts from the Hofmeister series and subsequently pelleted (Fig. 1a). Depolymerized tubulin and other soluble proteins remained in the supernatant, while intact axonemes and other insoluble proteins sedimented into the pellet. Based on this assay, we decided to test NaNO3 and KI, which both solubilize significant amounts of tubulin. We found that the more chaotropic salt, KI, leads to tubulin denaturation and no tubulin polymerization was observed after treatment with this salt. In contrast, NaNO3 has no significant effect on polymerization, and therefore was selected for our purifications.

**Figure 1.**
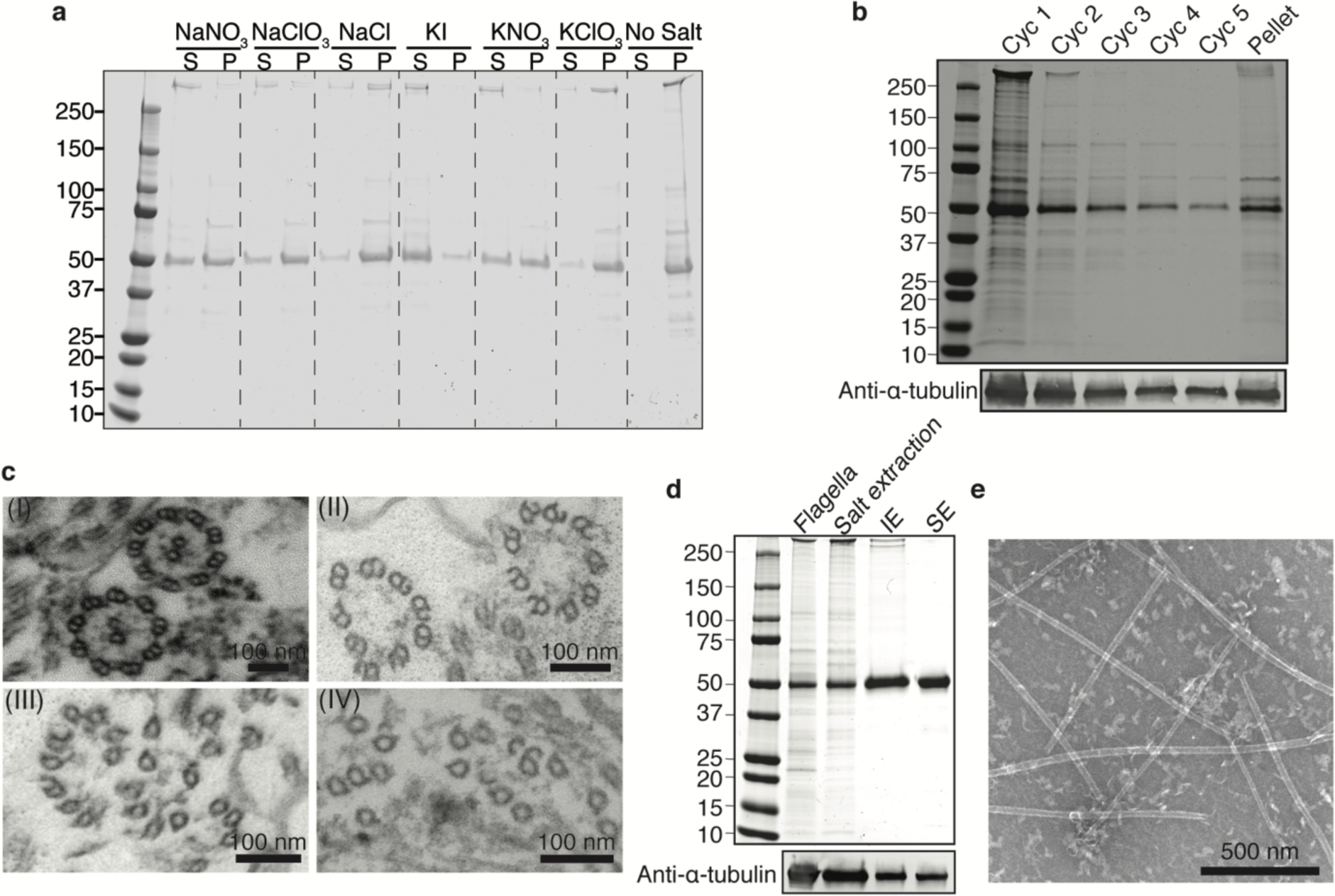
Purification of axonemal tubulin and MTs reconstitution. (**a**) SDS-PAGE of supernatant (**S**) and pellet (**P**) axoneme samples after salt extraction (500 mM). Supernatant contains soluble axonemal tubulin (50 kDa), while the pellet contains tubulin from insoluble axonemes. (**B**) SDS-PAGE of supernatant samples from five cycles of 500mM NaNO3 salt extraction, and the pellet at the end. (**C**) Transmission electron microscopy micrographs of (I) intact axoneme and axonemes after (II) 1 cycle (III) 2 cycles and (IV) 5 cycles of salt extraction with 500mM NaNO3. (**D**) SDS-PAGE of samples collected throughout the axonemal tubulin purification. From left to right: flagella lysate, supernatant of the axonemes after treating them with 500mM NaNO3 salt extraction, Hi-Trap Q (IE) elution fractions, size exclusion (SE) elution. (**E**) TEM micrograph of taxol stabilized reconstituted axonemal MTs.

We exploited the limited extraction of tubulin by NaNO3 to sequentially extract tubulins of different stabilities. The axonemes were subjected to five consecutive cycles of salt extraction (Fig. 1b). Interestingly, even after five cycles of extraction, with decreasing amount of tubulin extracted in each cycle, there was a significant amount of tubulin in the pellet. Intrigued by this observation, we examined the “9+2” architecture of the axoneme during the extraction process by transmission electron microscopy (TEM). After one cycle of extraction, the CP and a small part of the B-tubule were absent. After two cycles, another part of the B-tubule was missing, and after five cycles of salt extraction primarily A-tubules were observed in the pellet (Fig. 1c). Thus, different components of the axoneme structure exhibit various stabilities to salt extraction. The extracted tubulin was further purified by two chromatography steps - anion exchange and size exclusion - to separate tubulin from other microtubule-associated proteins (MAPs) (Fig. 1d). Finally, the functionality of the purified tubulin - the ability to polymerize into MTs - was confirmed by TEM (Fig. 1e).

### Axonemal MTs have dynamic properties distinct from brain MTs

We characterized the dynamic properties of the various axonemal tubulins using interference reflection microscopy (IRM) to perform label-free dynamic MT assays^27^ (Table 1). Purified tubulins (a mixture of CP and B-tubule tubulins) were assembled from GMP-CPP stabilized MT seeds polymerized from bovine brain tubulin and attached to the coverslip (Fig. 2a,b).

**Figure 2.**
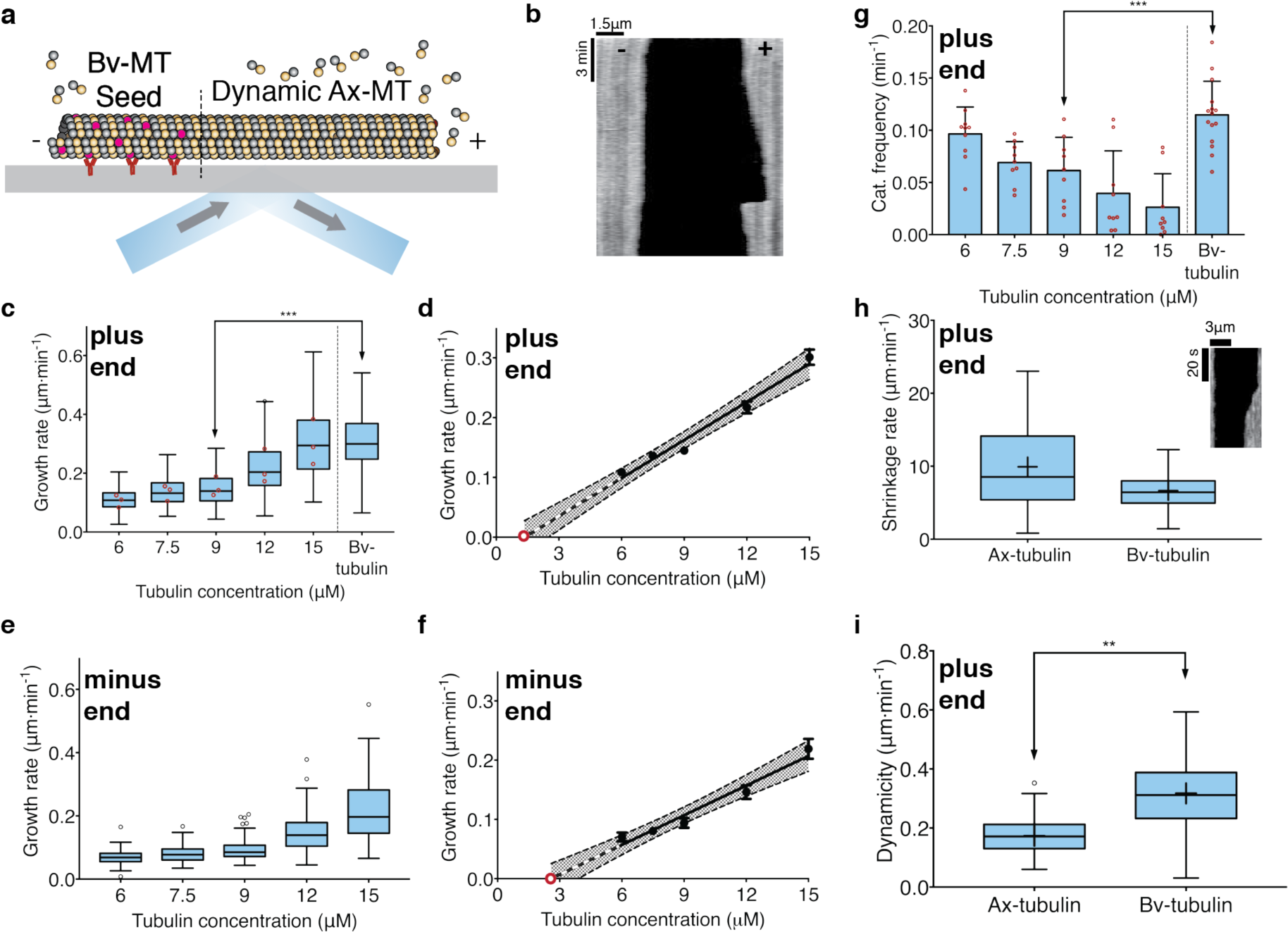
Polymerization dynamics of axonemal microtubules. (**a**) Schematic of the experimental design. Axonemal MTs grow from GMPCPP-stabilized TAMRA-labeled bovine MT seeds, which are immobilized to the coverslip with anti-TAMRA antibodies. (**b**) Representative kymograph of axonemal MT growth imaged using interference reflection microscopy. (**c**) Tukey plot showing growth rate of axonemal MTs in different tubulin concentrations, derived from three independent technical replicates for each of the three biological replicates. Each red circle marks the mean growth rate of a single biological replicate (*n*=275, 280, 268, 252, 265, 542; \*\*\**p*=0.0003) (**d**) The mean growth rates at different tubulin concentrations from panel (c) were fit by linear regression. No growth is observed below 6 µM and the *x*-intercept (red open circle) is 1.35 µM. Grey area denotes 95% confidence interval. Error bars represent S.E.M. (**e**) Tukey plot showing growth rate at the minus end of axonemal MTs in different tubulin concentrations (*n*=48, 79, 84, 132, 127). (**f)** The mean growth rates at different tubulin concentrations from panel (e) were fit by linear regression. The extrapolated concentration is 2.6 µM. Grey area denotes 95% confidence interval. (**g**) Bar plot showing catastrophe frequency of axonemal MTs at different tubulin concentration and bovine tubulin at 9 µM (*n*=10, 9, 9, 9, 9, 15; \*\*\**p*=0.0007). (**h**) Tukey plot showing the shrinkage rate of axonemal MTs and bovine MTs at 9 µm tubulin concentration (*n*=111, 211). Inset: kymograph showing a typical shrinkage event (**i**) Tukey plot showing the dynamicity of axonemal MTs and bovine MTs at 9 µM tubulin concentration (*n*=75, 75; \*\**p*=0.008).

**Table 1.** Rate constants for axonemal tubulin

|  | Plus end | Minus end |
| --- | --- | --- |
| **Slope** ($\mu\text{m}\cdot\mu\text{M}^{-1}\cdot\text{min}^{-1}$) | $0.021 \pm 0.001$ | $0.016 \pm 0.001$ |
| **K<sub>on</sub>** ($\mu\text{M}^{-1}\cdot\text{s}^{-1}$) | $0.54 \pm 0.03$ | $0.43 \pm 0.03$ |
| **y-intercept** ($\mu\text{m}\cdot\text{min}^{-1}$) | $-0.029 \pm 0.011$ | $-0.043 \pm 0.012$ |
| **k<sub>off</sub>** ($\text{s}^{-1}$) | $0.785 \pm 0.297$ | $1.164 \pm 0.325$ |
| **x-intercept** ($\mu\text{M}$) | $1.35 \pm 0.77$ | $2.58 \pm 0.67$ |

Values represent mean ± SEM

We first investigated how the growth rates depend on tubulin concentrations. The minimum concentration for elongation from the seeds was ~6 µM tubulin, similar to that in bovine brain tubulin. In comparison to bovine brain tubulin, however, the growth rate at the plus end of axonemal tubulin was significantly slower (0.15 ± 0.003 µm min^-1^ vs. 0.30 ± 0.004 µm min^-1^, all errors are SEM unless otherwise noted) (Fig. 2c). We then performed linear regression on the growth rate vs. tubulin concentration (Fig. 2d) and found that the extrapolated tubulin concentration for growth at the plus end (*x*-intercept) was 1.35 ± 0.77 µM, not significantly different from zero, and similar to bovine tubulin^28^. The slope of the axonemal tubulin curve corresponds to a second order association rate constant of 0.54 µM^-1^•s^-1^. This rate is significantly lower than previous measurements for brain tubulin^29^ (after correcting for temperature). Thus, axonemal MTs have slower growth rates than bovine tubulin.

To determine the differences between the two ends of the axonemal MTs we performed the same analysis for the minus end. As for the plus end, we found that the rate of polymerization was proportional to the tubulin concentration (Fig. 2e). Linear regression showed that the minimal concentration for elongation at the minus end is ~2 fold higher (2.58 ± 0.67 µM) compared to the plus end (Fig. 2f), and the slope of the curve corresponds to a second order association rate constant of 0.43 µM^-1^•s^-1^, only ~20% less than for the plus end. Thus, these results differ from brain tubulin which shows a 2-fold difference in the growth rates between the plus and minus ends^30^.

We then interrogated the catastrophe rate, which may play an important role in the axoneme length stability. The catastrophe rate at the plus end was measured for different tubulin concentrations and compared to bovine tubulin. The rate of catastrophe decreased as the concentration of tubulin increased, and compared to bovine tubulin, axonemal tubulin showed a 2-fold lower catastrophe rate (0.06 ± 0.01 min^-1^ vs. 0.12 ± 0.01 min^-1^) (Fig. 2g). Catastrophe events at the minus end, and rescue events at both ends were rarely detected in our assay, and for this reason, we omitted these parameters from the subsequent analysis.

To test whether the depolymerization (shrinkage) rate of axonemal tubulin is different than bovine tubulin, we followed it by IRM fast imaging (Fig. 2h). The shrinkage rate of axonemal tubulin is 1.5 times faster than that of bovine tubulin at the same concentration (9.9 ± 0.53 µm/min vs. 6.6 ± 0.15 µm/min).

Finally, we determined the dynamicity of axonemal and bovine tubulin from the dynamic assay (Fig. 2i). Dynamicity was calculated as previously reported by the sum of total growth and shrinkage lengths divided by the total time^31^. There was a 2-fold decrease in dynamicity of axonemal tubulin compared to bovine tubulin (0.17 ± 0.02 µm/sec vs. 0.32 ± 0.01 µm/sec). Thus, axonemal tubulin is less dynamic than bovine tubulin.

### Central-pair and B-tubule tubulins show no difference in dynamic properties despite different PTMs

We further investigated any possible dynamic differences between CP and doublet tubulins by purifying separate fractions of tubulin from the CP and the B-tubule. Because the CP MTs are known to be more labile than the doublet MTs^7^, we developed a protocol for differential extraction, in which one extraction at 330 mM NaNO3 was followed by four extractions at 500 mM NaNO3. Consequently, the first cycle included CP tubulin (TEM images confirmed lack of CP, Supplementary Fig. 1), while the other four cycles included mainly tubulin from the B-tubule, and the pellet comprised A-tubule MTs. Thus, we could soluble tubulin fractions highly enriched in either CP or B-tubule tubulin.

Previously it was reported that axonemal MTs from the CP, B-tubule and A-tubule have different PTMs. Hence, we analyzed the PTMs of the extracted tubulins in the different cycles by immunoblot (Fig. 3a). Anti-a-tubulin antibody verified that similar amounts of tubulin were found in the pellet and in the first two cycles of extraction, whereas in each of the following cycles the amount of extracted tubulin decreased. The extracted tubulin in the first cycle and the pellet had reduced polyglutamylation, while the tubulin from the other four cycles showed higher amounts of polyglutamylation. These results are consistent with previous study that showed that MTs in the CP and the A-tubule have a reduced amount of polyglutamylation compared to the B-tubule^32^. Tubulins from the CP, B-tubule and A-tubule were all acetylated, and contained a mixture of tyrosinated, detyrosinated and D2-tubulin. We then examined the glycylation of the axoneme. TAP952 and AXO49 antibodies, which are frequently used to detect glycylation in motile cilia, did not label any of the tubulins. However, Gly-Pep1, which labels glycylated tubulin in primary cilia^14^, labeled the tubulin in the CP and B-tubule, and to lesser extent tubulin from the A-tubule. Thus, our purification assay provides several new insights into the distribution of the PTMs that decorate the axonemal MTs (Fig. 3b).

**Figure 3.**
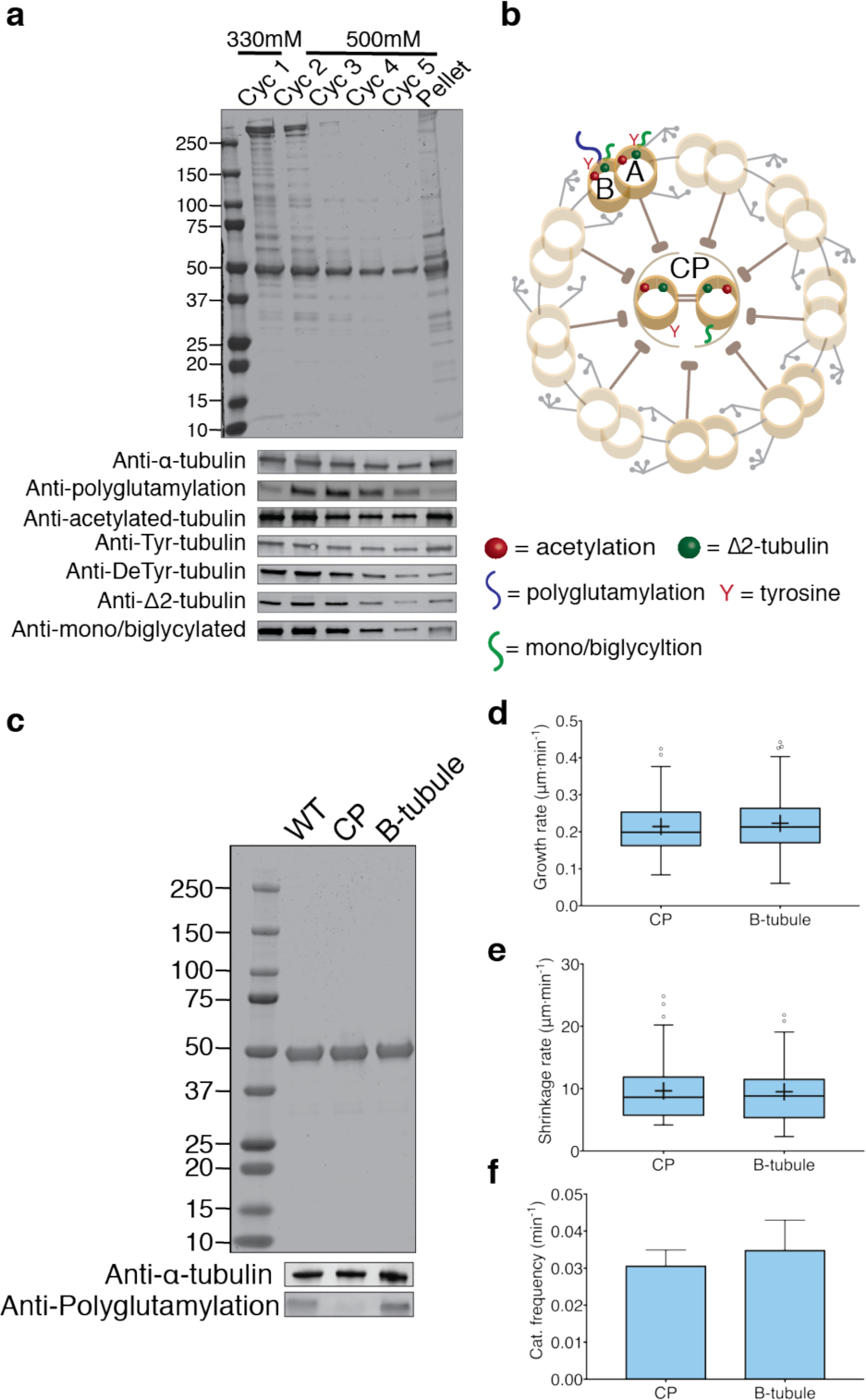
Purification, post-translational modifications analysis and dynamics of central pair and B-tubule tubulins. (**a**) SDS-PAGE and immunoblots of samples from salt-extracted cycles. Lower NaNO3 concentration in the first cycle allows mainly depolymerization of CP. (**b**) Schematic of the different PTMs on the axoneme structure. (**c**) SDS-PAGE and immunoblot of purified axonemal tubulin from whole axoneme, CP-enriched fraction and B-tubule. (**d**) Tukey plots showing growth rate and (**e**) shrinkage rate of tubulin from the CP and B-tubule at 9 µM tubulin concentration. (**f**) Average catastrophe frequency of axonemal MTs polymerized from CP and B-tubule at 9 µM tubulin concentration, derived from three independent experiments (*n*>120). Error bars represent S.D.

To determine whether these PTMs have an effect on dynamics, we further purified the tubulins from the CP, and the B-tubule (Fig. 3c). No significant differences were observed for growth rate (Fig. 3d), shrinkage rate (Fig. 3e), or catastrophe frequency (Fig. 3f). These results were supported by the characterization of the dynamic properties of two other *C. reinhardtii* mutants: (1) *pf18* that completely lacks the CP so that most of the purified axonemal tubulin consists of polyglutamylated tubulin. (2) *tpg1* that lacks tubulin tyrosine ligase like 9 (TTLL9) and has a reduced amount of polyglutamylation^19^. In both strains no significant differences were observed in their dynamic properties (Supplementary Fig. 2). Thus, our experiments with a single isotype tubulin indicate that there are no detectable differences in the dynamic properties between CP and B-tubule tubulins.

### Axonemal MTs form curved tip structures

During the analysis of the reconstituted axonemal MTs we observed a unique phenomenon. At tubulin concentrations above 9 µM, 5-10% of the growing MTs formed curved tip structures, not observed for brain MTs. The tips were dim compared to the MT shaft, indicating a lower number of PFs (Fig. 4a), and showed a range of lengths and curvatures (Fig. 4b). The mean tip length of ~1.5 µm was independent of tubulin concentration, while the tip curvature increased from 2.0 ± 0.8 µm^-1^ at 9 µM to 2.7 ± 1.1 µm^-1^ at 15 µM (mean ± SD) (Table 2 and Supplementary Fig. 3a-b). The distal tip curvature showed a small but significant decrease with tip length (Supplementary Fig. 3c). We further examined the tips of axonemal MTs by TEM and found open sheets at the MT ends (Figure 4c). We therefore propose that the tip structures are unclosed sheets comprising several PFs.

**Figure 4.**
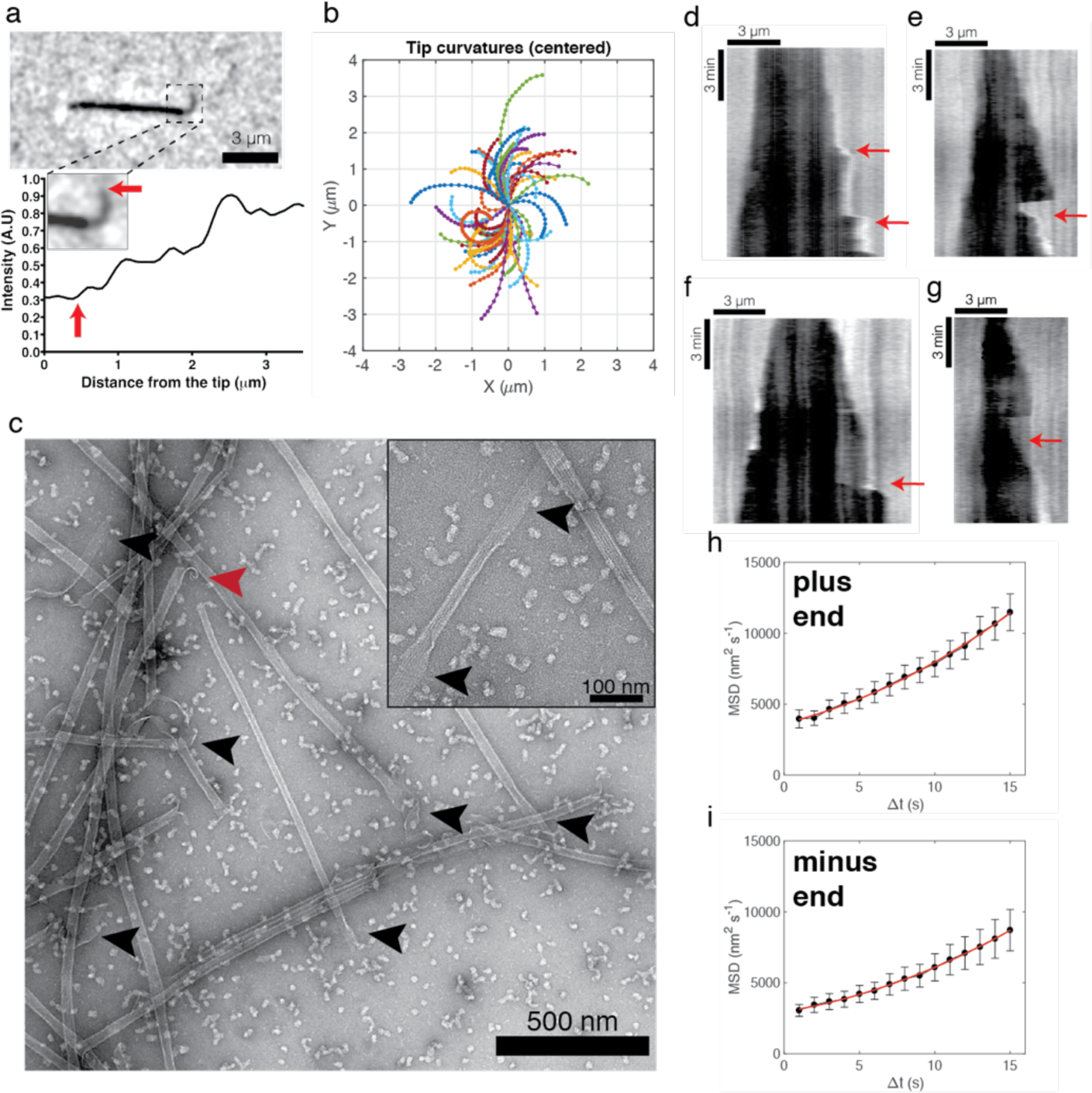
Growth of curved protofilaments polymerized from axonemal tubulins. (**a**) An image of curved PFs (dim region) growing from a MT (top). Intensity profile of the MT tip show stepwise increases (bottom). Red arrow indicates the tip. (**b**) Traces of individual curved PFs. (**c**) TEM image of growing MTs with open sheets at the ends (indicated in black arrows), and shrinking MT with “ram’s horns” (red arrow). Inset: high magnification of MT with open sheets at its ends. (**d**) Kymograph showing fast growth phases. (**e**) Kymograph showing fast growth phase after catastrophe event. (**f**) Kymograph showing the growth of a few PFs (dim region) and the growth of additional PFs alongside them. (**g**) Kymograph showing stable PFs after catastrophe event. Red arrows indicate each event. (**h**+**i**) MSD against time. A quadratic fit yielded diffusion coefficient and velocity of plus end and minus end.

**Table 2.** Structural properties of axonemal curved tips

| | 9 $\mu\text{M}$ | 12 $\mu\text{M}$ | 15 $\mu\text{M}$ |
| --- | --- | --- | --- |
| **Mean length** ($\mu\text{m}$) | $1.53 \pm 0.07$ | $1.36 \pm 0.07$ | $1.67 \pm 0.08$ |
| **Min length** ($\mu\text{m}$) | 0.61 | 0.65 | 0.65 |
| **Max length** ($\mu\text{m}$) | 2.87 | 2.69 | 3.32 |
| **SD** ($\mu\text{m}$) | 0.48 | 0.51 | 0.55 |
| **Mean Curvature** ($\mu\text{m}^{-1}$) | $1.95 \pm 0.12$ | $2.25 \pm 0.13$ | $2.71 \pm 0.17$ |
| **SD** ($\mu\text{m}^{-1}$) | 0.77 | 0.92 | 1.13 |

Values represent mean ± SD

This hypothesis is consistent with several unusual growth features seen for MTs with tip structures. First, the tips darkened as they straightened (Supplementary video 1), consistent with a transition from a curved sheet to a straight MT. Second, we observed fast growth phases of the MTs (Fig. 4d, Supplementary video 2), including after catastrophe events (Fig. 4e, Supplementary video 3). These observations are consistent with incomplete tip structures providing a template for rapid MT growth. Third, we observed rapid growth along a dim tip structures (Fig. 4f, Supplementary video 4). Fourth, occasionally we identified dim regions of the MTs after catastrophe events, suggesting that only some PFs underwent catastrophe (Fig. 4g, Supplementary video 5). This observation was confirmed by fast imaging of MTs that form a dim curved tip after a catastrophe event (Supplementary videos 6 and 7). Taken together, these observations suggest that PF sheets provide stable structures that allows fast templated growth.

Though the curved tips were only observed in 5-10% of MTs imaged by IRM, we wondered whether they might be present in a larger fraction of MTs, but not seen due to their shorter lengths. One characteristic of growth in the presence of curved tips is the variability of growth velocities. To determine whether variable growth is a property of all the axonemal MTs, we tracked the position of MT ends without observed PFs, and performed a mean-squared displacement (MSD) analysis in which the average of the square of the displacement in a time interval is plotted against the time interval (Fig. 4hi). By fitting our results to a quadratic equation, we calculated the average growth rate (*V*) and the diffusion coefficient (*D*), which is a measure of the variability of the growth rate. The diffusion coefficient at the plus end is much larger than expected if the only source of growth variability is the random arrival of tubulin dimers that bind irreversibly at the end (Table 3). Thus, all MTs display growth variability, even those that do not display tip structures. Previously, such high diffusion coefficients have been interpreted as arising from reversible binding of tubulin dimers at the growing tip, so that a majority of the tubulins that bind also unbind^29^. However, as we argue in the Discussion, the PF templated growth may provide as an alternative interpretation.

**Table 3.** Estimated rate constants from MSD analysis

|  | Plus end | Minus end |
| --- | --- | --- |
| *Number of MTs (n)* | 45 | 43 |
| *Diffusion coefficient (D, nm<sup>2</sup> s<sup>-1</sup>)* | $130.26 \pm 15.30$ | $87.50 \pm 10.39$ |
| *Growth velocity (V, nm s<sup>-1</sup>)* | $4.14 \pm 0.22$ | $3.72 \pm 0.17$ |
| *Expected Diffusion coefficient\* (D\*, nm<sup>2</sup> s<sup>-1</sup>)* | $2.55 \pm 0.13$ | $2.28 \pm 0.10$ |
| **k<sub>on</sub>[Tub]** ($\text{s}^{-1}$) | $347.76 \pm 40.41$ | $234.36 \pm 27.45$ |
| **k<sub>on</sub>** ($\mu\text{M}^{-1}\text{s}^{-1}$) | $38.64 \pm 4.49$ | $26.04 \pm 3.05$ |
| **k<sub>off</sub>** ($\text{s}^{-1}$) | $341.03 \pm 40.46$ | $228.33 \pm 27.47$ |
| **$\sigma^2$** (nm<sup>2</sup>) | $60.33 \pm 0.88$ | $54.32 \pm 0.66$ |

Values represent mean ± SEM
\*Diffusion coefficient assuming random and irreversible tubulin dimer binding, $D^* = V \times 8$ nm/13 PFs

## Discussion

### Axonemal tubulin has distinct dynamic properties that are not affected by PTMs

To understand the role of MTs in the length stability of axonemes, we studied axonemal tubulin *in vitro*. To extract both CP and doublet tubulins, we used the Hofmeister series to destabilize the axonemes, and by adjusting the salt concentration of NaNO3 we were able to purify, for the first time, single isotype functional tubulins from the CP and the B-tubule. We found that axonemal tubulin shows a 2-fold decrease in dynamicity compared to bovine tubulin, which is reflected by a slower growth rate and lower catastrophe frequency. Thus, the lower dynamicity is in accord with the higher length stability of the axoneme structure.

Axonemal tubulin still undergoes dynamic instability, however, and we asked whether PTMs directly contribute to the axoneme length stability. Previous studies, which were based on mammalian brain tubulin, involved several tubulin isotypes and consequently the effect of PTMs on the dynamic properties has remained elusive. However, *C. reinhardtii* has a single tubulin isotype and therefore we can isolate this effect. Thus, we first characterized the PTMs on the different axoneme components (CP, B-tubule and A-tubule). We found that polyglycylation, which was suggested to be involved in axoneme length stability but was never identified in *C. reinhardtii*, is indeed decorating all axonemal MTs. We also found, in contrast to a previous study^33^, that both CP and A-tubule contain detyrosinated- and D2-tubulin. Thus, all the axoneme components consist of tyrosinated-, detyrosinated- and D2-tubulin. Although there is no known function for these PTMs in cilia, it was recently suggested that detyrosination of MTs regulates cardiac mechanotransduction^34^, and therefore might be playing similar role also in cilia. Finally, in agreement with a previous report^32^, we found that polyglutamylation, which was suggested to regulate flagellar motility^19,21,35,36^, predominantly decorates the B-tubule. However, our assays with tubulins from the CP and B-tubule, as well as two different *C. reinhardtii* mutants, reveal that polyglutamylation has no significant effect on the dynamic properties of MTs. Whether other PTMs or combination of different PTMs affect the dynamic properties of tubulin will require further study.

### High stability of the PFs provides a mechanism for MT recovery

During the characterization of the axonemal MTs, we discovered that at high tubulin concentrations (> 9 µM) some of the growing axonemal MTs have relatively dim, curved tips at their ends. We do not think that the tip structures are caused by contamination of MAPs because our purification scheme includes a stringent salt extraction followed by two chromatography steps, anion exchange and size exclusion. This was further supported by mass spectrometry analysis, which failed to identify any known MAPs (Supplementary Table S1). The lower intensity and high curvature, distinguish these tip structures from the cylindrical structure of complete MTs, and suggest that they are formed by unclosed protofilament sheets, which we also observed in TEM, and others have observed in cryo-EM^37-39^. Our failure to observe them for bovine MTs suggests that axonemal PFs maybe more stable than those of brain tubulin.

Although the tip structures were detected only on some of the MTs and at high concentrations, we speculate that it may be a general feature that occurs on most MTs, even at lower concentrations. However, due to the short tip lengths and/or low number of PFs we cannot detect them in our assays. This hypothesis was supported by our MSD analysis that showed high growth variability of the axonemal MTs.

High PF stability may account for several unusual features of growth. For example, fast growth phases are observed, despite the low average dynamicity of the MTs. Moreover, the PFs show unique stability to catastrophe events, which is reflected by PFs that do not depolymerize after these events, and provide a template for newly polymerizing PFs. This provides a mechanism for fast recovery of the MTs from catastrophe events. While we observe these phenomena only in axonemal MTs, this type of behavior might happen to a lesser extent in other tubulins with much shorter tip structures as observed in some EM studies, and could explain the rapid fluctuations in their growth rates^29,40,41^.

What are the molecular differences between axonemal MTs and other MTs? The answer to this question might be related to differences in the lateral and longitudinal interactions. Previous studies showed that Lys-60 stabilizes the lateral contacts between adjacent subunits^42,43^. However, in *C. reinhardtii* Lys-60 is substituted for Arg in b-tubulin. This can provide a longer side chain, and tolerance to structural changes that accompany the MT before catastrophe events^43^ (Supplementary Fig. 4). Intriguingly, we found that Arg-60 is common among ciliates, whereas in other species that are commonly used to study MTs, Lys-60 is highly conserved (Supplementary Fig. 5). Another substitution is Tyr-224 that is located at the E-site and interacts with the nucleobase and with Asp-179, which takes an active role in the formation of longitudinal interactions^44^. In *C. reinhardtii* Tyr-224 is substituted for a Phe residue, which is missing the hydroxyl group and therefore cannot interact with Asp-179 (Supplementary Fig. 6). This frees Asp-179 to promote longitudinal contacts. Another role of Tyr-224 is in the positioning of the GTP in the E-site. However, the interaction of Phe with the nucleobase might lead to different positioning and consequently affecting GTP hydrolysis. As in the other case, Tyr-224 is highly conserved between other species that are used to study MTs (Supplementary Fig. 7), while Phe-224 is found only in plants^45^. Thus, we propose a structural basis for the unique properties of axonemal tubulin, which could be tested by additional experiments.

### The growth of axonemal MTs suggests that GTP-tubulin has a curved conformation

A long-standing question concerns the mechanism of MT assembly by GTP-tubulin. Early observations of straight PFs at the MT tips during polymerization, and curved PFs during depolymerization, paved the way to a canonical model^46^. According to this model GTP-tubulin forms a straight conformation that is more compatible with the MT lattice, and thus promotes the polymerization of MTs. The delayed hydrolysis of GTP to GDP after polymerization leads to strained GDP-tubulin configuration that is forced to adopt an energetically unfavorable straight conformation within the MT lattice. This strain energy is released by catastrophe events that allows the GDP-tubulin to assume its preferred curved conformation^47^.

In contradiction to the canonical model, an increasing number of studies support the notion that tubulin has a curved conformation independent of its nucleotide state^48^. This is based on the crystal structures of tubulin^44,49-51^, and cryo-EM analyses that showed slightly curved PFs at the end of growing MTs^37-39,52,53^. These studies were further supported by biochemical experiments^49-51,54,55^. However, due to technical reasons, such as using unpolymerized tubulin, and the absence of curved tips in dynamic assays, the curvature of GTP-tubulin is a matter of debate.

Our dynamic assays provide new insight into MT tip growth (Figure 5), since all previous studies were based on static images. Our assays indicate that growing MTs generate curved tip structures; if the tubulin in these structures is in the GTP state, as had been argued by Odde et al.^29^, then our observations support a curved GTP-tubulin configuration. The radius of curvature at the very ends of the tips is ~400 nm, an order of magnitude larger the tightly curved “ram’s horns” observed in shrinking MTs that are thought to represent the GDP state^53^. These values are also much larger than the values previously reported by cryo-EM studies^37^-39, suggesting that the curved tip structure in our assays include higher number PFs. Future studies will include cryo-EM characterization that will a provide better resolution to axonemal MT tips.

**Figure 5.**
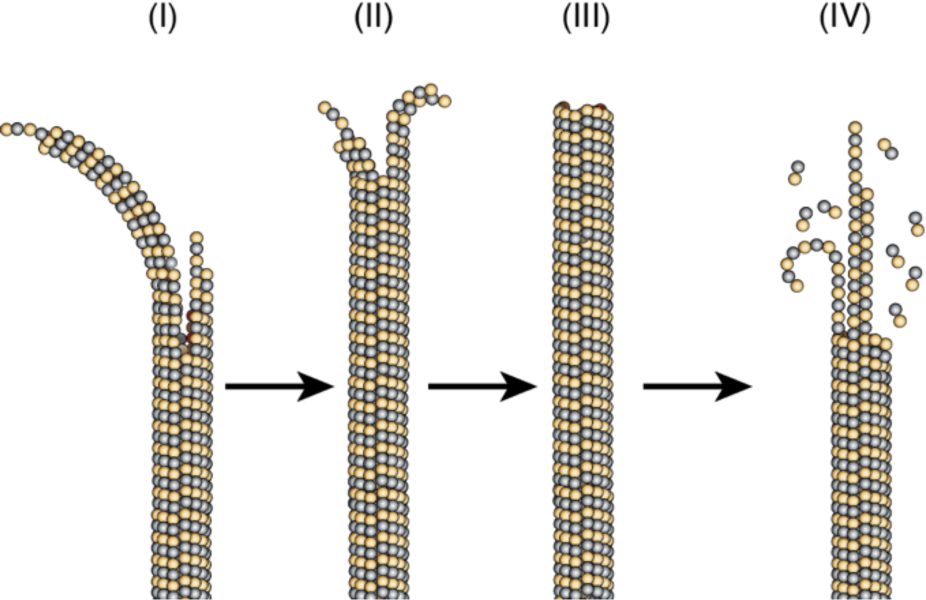
Model for growth and shrinkage for axonemal tubulin. (**I)** MTs polymerize by incorporation of GTP-tubulin at their tip. During this process PFs form curved tip structures with radius of curvature of ~400 nm. (**II**) These curved PFs provide a template for polymerization of new PFs that catch up with the previous PFs, and thus leading to conformational change that will decrease the curvature of the tip. (**III**) The straight and full MT will be formed by the polymerization of 13 PFs. (**IV**) Upon catastrophe, a few PFs remain behind and provide a template for new PFs, while other can form “rams horns” structure that have much smaller radius of curvature.

In conclusion, *C. reinhardtii* provides a unique platform to study MTs structural and biophysical properties. Our results show that the dynamic properties of axonemal tubulin together with its structural properties provide two different mechanisms that can support the high length stability of the axoneme structure. The single tubulin isotype in *C. reinhardtii* allows us to study the effect of PTMs without the complexity of isotypic heterogeneity. Thus, we can conclude that polyglutamylation has no effect on the dynamic properties of *Chlamydomonas* axonemal tubulins. Finally, axonemal tubulin also provides a new insight into the growth of MTs and specifically the conformation of GTP-tubulin.

## Acknowledgments

We thank Dr. Xinran Liu from the core center for cellular and molecular imaging at Yale for sample processing, sectioning and image acquisition, and Dr. Joel Nott from the protein facility at Iowa State University for the mass-spectrometry analysis. Special thanks to Dr. Sabyasachi Sutradhar for the great help with MATLAB and all The Howard lab members for fruitful discussion. Research reported in this publication was supported by a Cross-Disciplinary Fellowship from Human Frontier Science Program (LT000919/2015-C) and European Molecular Biology Organization long-term fellowship (ALTF 1424-2014) to RO, and National Institute of General Medicine Sciences of the National Institutes of Health under award number R01GM110386 to JH.

## Author contributions

R.O designed and performed all experiments. R.O and J.H analyzed data and wrote the paper.

## Materials and methods

### *Chlamydomonas* strains and culture conditions

*C. reinhardtii* wild-type strain 137c (CC-125), pf18 (CC-1036) and tpg1(CC5245) were obtained from the *Chlamydomonas* Resource Center (St. Paul, MN). Cells were cultured as previously described^56^. Briefly, strains were maintained on plates of TAP medium (20 mM Tris HCl, 7 mM NH4Cl, 0.40 mM MgSO4, 0.34 mM CaCl2, 2.5 mM Na3PO4, and 1000× diluted Hutner’s trace elements, titrated to pH 7.0 with glacial acetic acid)^57^ containing 1.5% agar. For axonemal tubulin purification, cells were resuspended in liquid TAP media and grown with continuous aeration and illumination at 21°C for ~7 d.

### Bovine tubulin purification and labeling

Bovine brain tubulin was purified by two cycles of polymerization-depolymerization and stored in 1ÍBRB80 (80mM PIPES, 1mM MgCl2, 1mM EGTA, pH 6.8)^58^. Protein concentration was determined by NanoDrop 2000 spectrophotometer (Thermo Fisher Scientific, USA). Purified protein was snap-frozen using liquid nitrogen and stored at −80°C until use. Tetramethylrhodaminelabeled tubulin was labeled using 5(6)-TAMRA, succinimidyl ester (Invitrogen, USA) as described previously^59^.

### Axonemal tubulin purification

*C. reinhardtii* cells were harvested and axonemes were isolated by standard methods^60^. Cells were deflagellated using 4.2 mM dibucane-HCl, and flagella were demembranated by 1% IGEPAL CA-630. Tubulin was then extracted by cycles of NaNO3 (330 mM/500 mM) salt extraction solution and centrifugation at 125,000g. Supernatant was collected and further purified by 5 ml HiTrap Q HP column (GE, USA), followed by size exclusion Superdex 200 increase 10/300 GL column (GE, USA). The purified axonemal tubulin protein was eluted in 1ÍBRB80 and samples were snap-frozen using liquid nitrogen and stored at −80°C until use. Protein concentration was determined by NanoDrop 2000 spectrophotometer.

### SDS-PAGE and Immunoblot

SDS-PAGE was performed using 4-15% Mini-PROTEAN TGX precast protein gels that were stained in Coomassie blue. For Western blot analysis, the following primary antibodies were used: Gly-Pep1 anti-mono/biglcylated tubulin (1:10,000; AdipoGen)^14^ anti-acetylated a-tubulin (clone 6-11B-1; 1:50,000; MilliporeSigma)^61^; anti-polyglutamylation (clone GT335; 1:1,000; AdipoGen)^62^; anti-detyrosinated tubulin (1:1,500); anti-D2 tubulin (1:1,000; MilliporeSigma) anti a-tubulin (clone B-5-1-2; 1:4,000; MilliporeSigma) anti-tyrosinated tubulin (1:1000; MilliporeSigma).

### Electron microscopy

For transmission electron micrscopy (TEM), axonemes were fixed (0.1 M sodium cacodylate buffer, 2.5% glutaraldehyde, 2% paraformaldehyde, pH 7.4) for 30 min at room temperature, followed by 30 min at 4°C. Samples were then rinsed 3 times in 0.1 M sodium cacodylate buffer, trimmed and postfixed in 1% osmium tetroxide for 1 h, en bloc stained in 2% uranyl acetate in maleate buffer at pH 5.2 for a another hour, rinsed and then dehydrated in an ethanol series, infiltrated with resin (Embed812 Electron Microscopy Science), and baked overnight at 60°C. Hardened blocks were cut using a Leica UltraCut UC7, and 60-nm sections were collected on formvar/carbon-coated nickel grids and stained using 2% uranyl acetate and lead citrate. Samples were imaged using a FEI Tecnai Biotwin transmission electron microscope. Images were taken using Morada CCD and iTEM (Olympus) software. For negative staining, 5 μl samples were deposited on a glow-discharged formvar/carbon-coated copper grid (Electron Microscopy Sciences) and stained with 2% uranyl acetate. Imaging was performed on a JEOL JEM-1400 Plus microscope.

### Microscopy assay and imaging conditions

For dynamic MT assays, axonemal tubulins extensions were grown from TAMRA-labeled GMPCPP-stabilized bovine MT seeds and imaged by Interference Reflection Microscopy (IRM)^27^. Coverslips (Marienfeld, Germany) were cleaned using piranha solution and rendered hydrophobic by silinization^59^. Experiments were performed in a flow channel formed by two Parafilm strips between 18×18 mm^2^ and 22×22 mm^2^ coverslips.

MTs growth was initiated by perfusing polymerization solution BRB80 supplemented with oxygen scavengers (20mM glucose, 20 µg/ml glucose oxidase, 8 µg/ml catalase, 0.1 mg/ml casein, 1mM dithiothreitol and 1mM GTP and tubulin at different concentrations. Dynamic assays were performed at 28°C and imaged at frame rates of 0.2 fps for growth rate analysis and 10 fps for shrinkage rate analysis.

### Identification of plus and minus ends

Because the growth rates at the plus and minus ends of axonemal MTs are sometimes indistinguishable, we used polarity marked MT seeds to identify the ends of the MTs. On average, the plus ends grew faster than the minus ends (Supplementary Figure S8), as expected. Furthermore, 76 out of 85 of these polarity marked MTs grew faster at their plus ends than their minus. Therefore, in the IRM measurements, we defined the MTs plus end as the end that grew more. The error in the assignment of MT polarity is therefore approximately 10%.

### Dual-Labeled GMPCPP stabilized MT seeds

GMPCPP stabilized seeds were polymerized from 2 µM TAMRA-labeled bovine tubulin for 20 min at 37°C. Then, the MTs were centrifuged for 5 min at 20 psi in an Airfuge, and resuspended in warm polymerization solution (BRB80, 1 mM GMPCPP, 1mM MgCl2, 0.6 µM Alexa488-labeled bovine tubulin). The solution was incubated for 30 min at 37°C, and then centrifuged for 5 mins in an Airfuge and resuspended in 200 µL warm BRB80.

### Image analysis

Image analysis was performed by creating kymographs of individual MT seeds using ImageJ analysis software (National Institutes of Health). Growth rates were calculated from the slopes of the kymographs.

### Statistics

Data plotting and curve fitting was performed with Prism 7 (Graph-Pad). Evaluations of statistical significance are described in the respective figure legends. Biological replicates are measurements of tubulin from different purifications. Technical replicates are measurements of tubulin from the same purification that were done in different days. All experiments were performed at least three times.

### Sequences

Table 4 presents the sequences that were used in this study. Tubulin sequences have been aligned with Geneious software. Amino acid homologies are marked in dots.

**Table 4.**
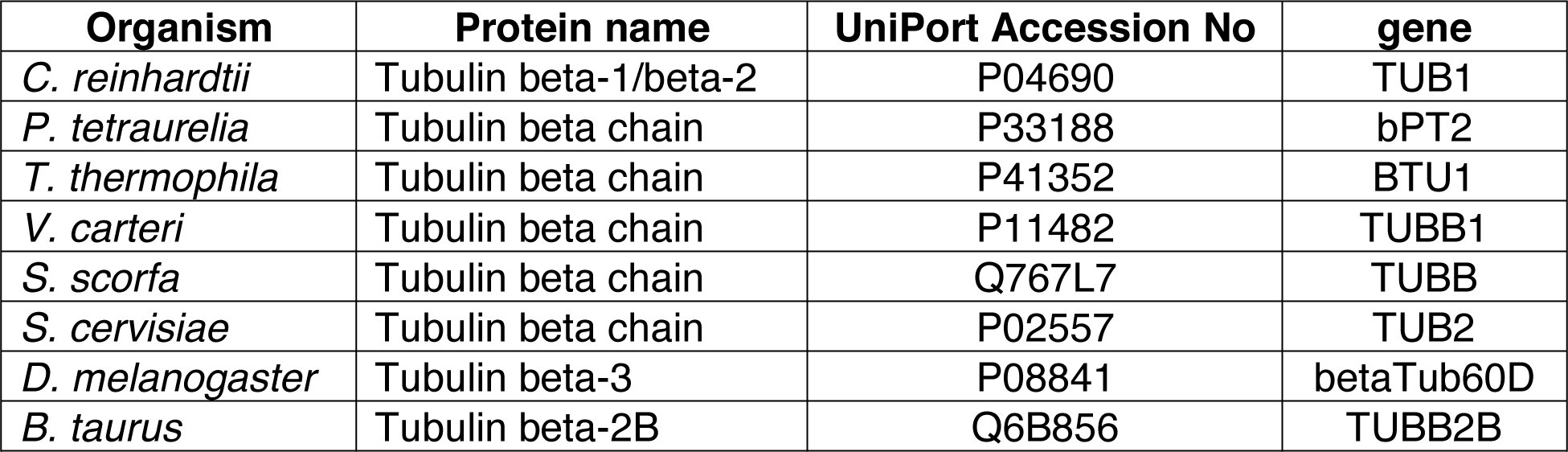
Tubulin sequences that were in this study

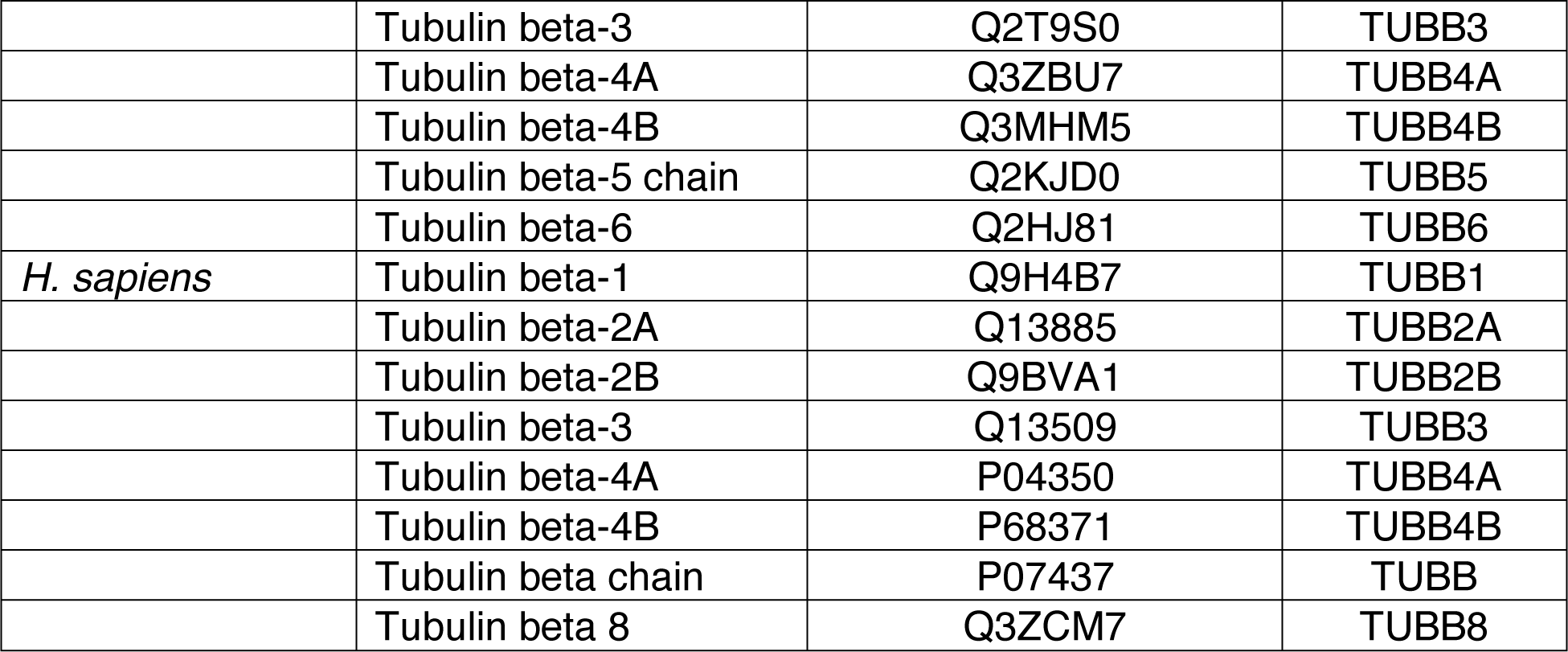

### MSD analysis

To determine the diffusion coefficient and net growth rate of axonemal tubulin, the end position of the MTs was tracked using FIESTA^63^. Data was then analyzed using the MSD analyzer package^64^, and values were calculated by quadratic fitting to the first 15 s.

### PFs analysis

The lengths of the MT tips were measured manually using ImageJ software. Start point of the measurement was determined by reduced intensity of the MT. Curvature measurements were performed in MATLAB by fitting a circle to the last 0.6 μm of the tip.

---

**Supplementary Figure 1.**
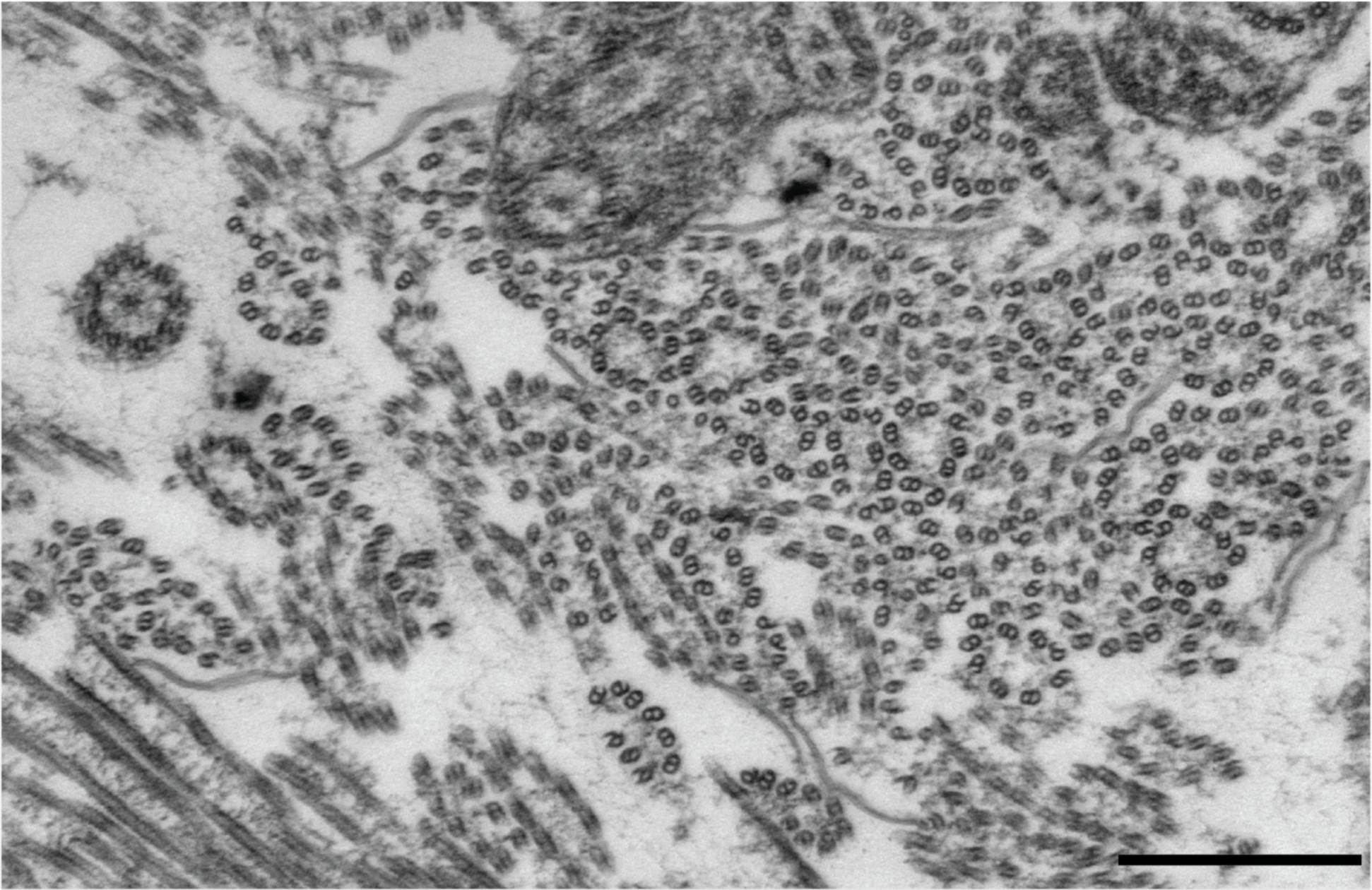
Differential extraction of axonemal tubulin. TEM micrograph of axonemes treated with 330mM NaNO3. scale bar: 500 nm.

**Supplementary Figure 2.**
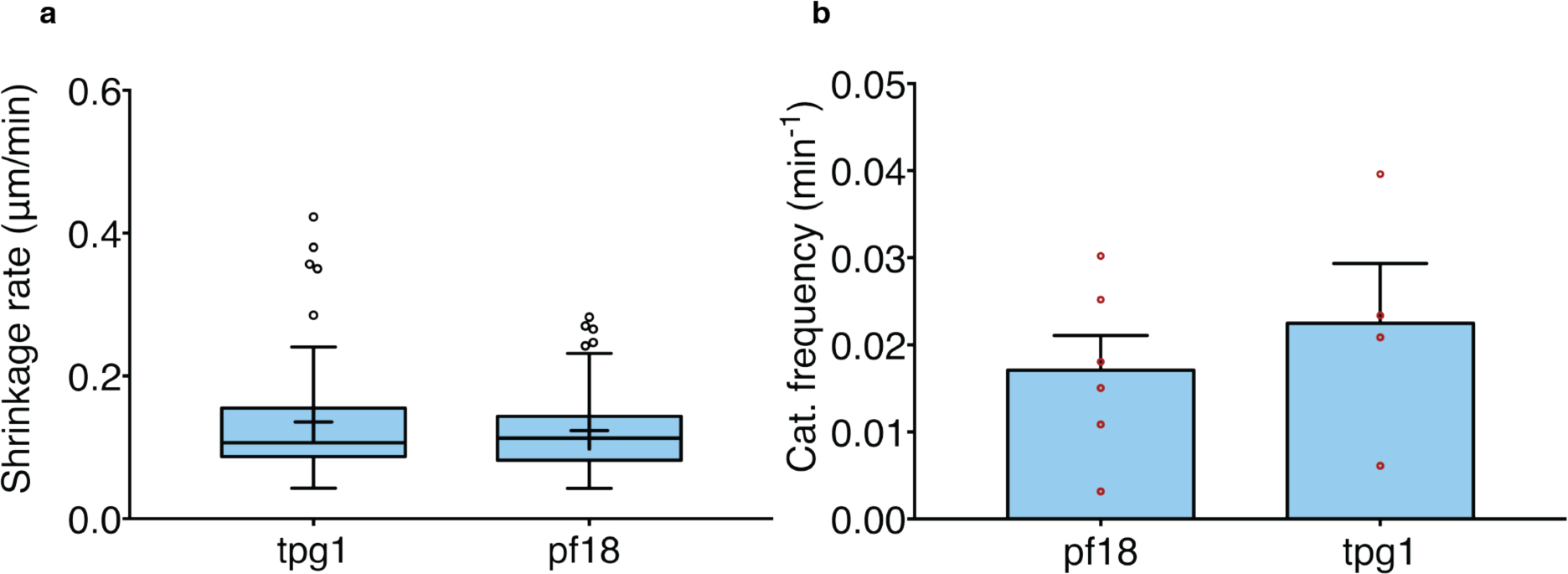
Dynamic properties of *C reinhardtii* mutants. (a) Tukey plot showing growth rate of axonemal MTs at tubulin concentrations of 9 µM. (*n* = 82, 201). (b) Catastrophe frequency of axonemal MTs at a tubulin concentration of 9 µM. Each red circle marks the catastrophe frequency of a single biological replicate. Error bars represent S.D.

**Supplementary Figure 3.**
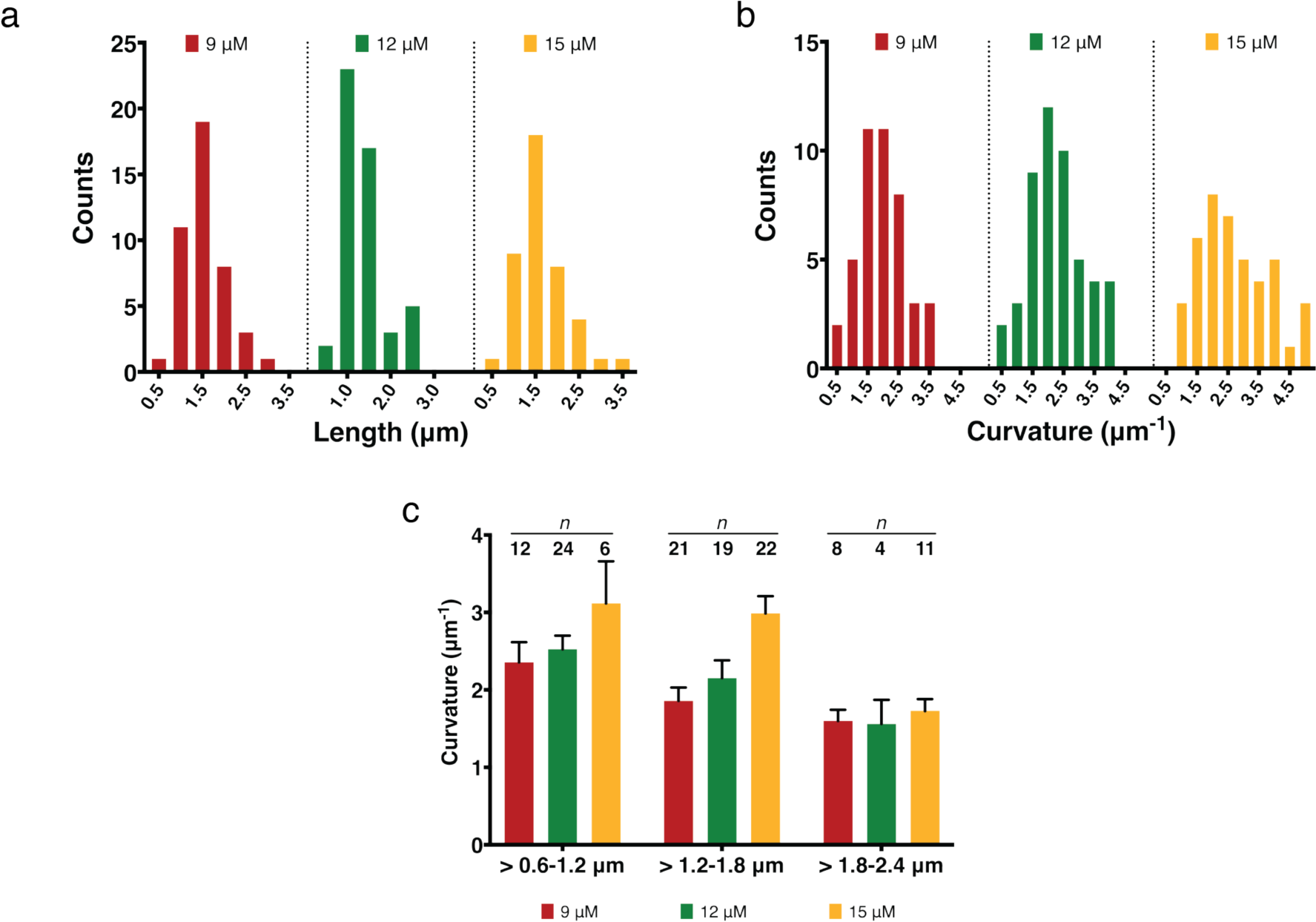
Growth of curved PFs polymerized from axonemal tubulins. (**a**) Histogram showing distributions of lengths of curved regions for different tubulin concentrations. (**b**) Histogram showing distributions of curvature for different tubulin concentrations. (**c**) Mean curvature for each tubulin concentration and for different tip lengths show linear trend.

**Table S1.** Mass spectrometry analysis of axonemal tubulin

| Accession | Description | Score | Coverage % | # Unique Peptides | Total Peptides | Protein Length | MW [kDa] |
| --- | --- | --- | --- | --- | --- | --- | --- |
| Q540H1 | Tubulin alpha chain OS=Chlamydomonas reinhardtii OX=3055 GN=TUA1 PE=3 SV=1 | 90513 | 82 | 33 | 1892 | 451 | 49.6 |
| A8IXZ0 | Tubulin beta chain OS=Chlamydomonas reinhardtii OX=3055 GN=TUB1 PE=3 SV=1 | 79389 | 84 | 37 | 2583 | 443 | 49.6 |
| NP_463397.1 | PTS system mannose-specific transporter subunit IID [Salmonella enterica subsp. enterica serovar Typhimurium str. LT2] | 91 | 3 | 1 | 5 | 278 | 30.9 |
| P02769 | Serum albumin OS=Bos taurus GN=ALB PE=1 SV=4 | 34 | 2 | 1 | 1 | 607 | 69.2 |
| NP_461896.1 | L-fuculokinase [Salmonella enterica subsp. enterica serovar Typhimurium str. LT2] | 32 | 2 | 1 | 2 | 472 | 51.6 |
| 809009895 | membrane protein [Erwinia tracheiphila] | 24 | 2 | 1 | 5 | 338 | 35.8 |
| NP_462668.2 | PTS system mannose-specific transporter subunit IID [Salmonella enterica subsp. enterica serovar Typhimurium str. LT2] | 22 | 4 | 1 | 2 | 285 | 30.7 |
| NP_462444.1 | gluconate operon transcriptional repressor [Salmonella enterica subsp. enterica serovar Typhimurium str. LT2] | 20 | 2 | 1 | 3 | 331 | 36.5 |

**Supplementary Figure 4.**
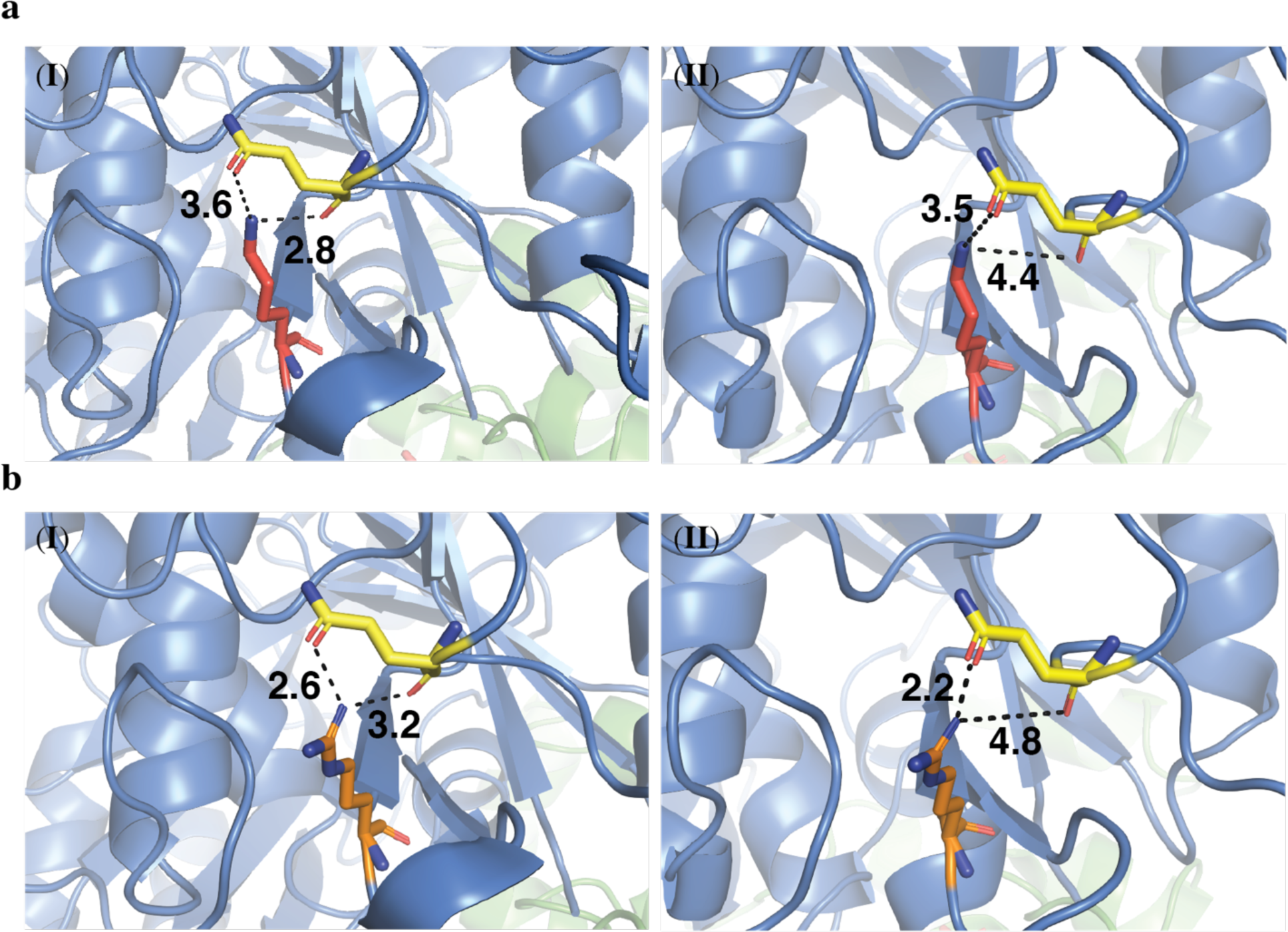
(**a**) Lys-60 (red) in the H1’-S2 loop interacts with Gln-280 (yellow) and stabilizes the lateral contacts between neighboring subunits at the GDP-Pi state (PDB: 6EVX) (I), Whereas in the GDP state this interaction is no longer involved due to the long distance (PDB: 6EVZ) (II). (**b**) Substitution of Lys-60 to Arg (orange) might lead to more stable interactions with Gln-280 in the GDP-Pi state due to the short distance (I) or even in the GDP state (II). Each substitution was plotted on the solved protein structure for the GDP•Pi-DCX-MT (PDB: 6EVX) using PyMOL software (1.1r1, http://www.pymol.org/).

**Supplementary Figure 5.**
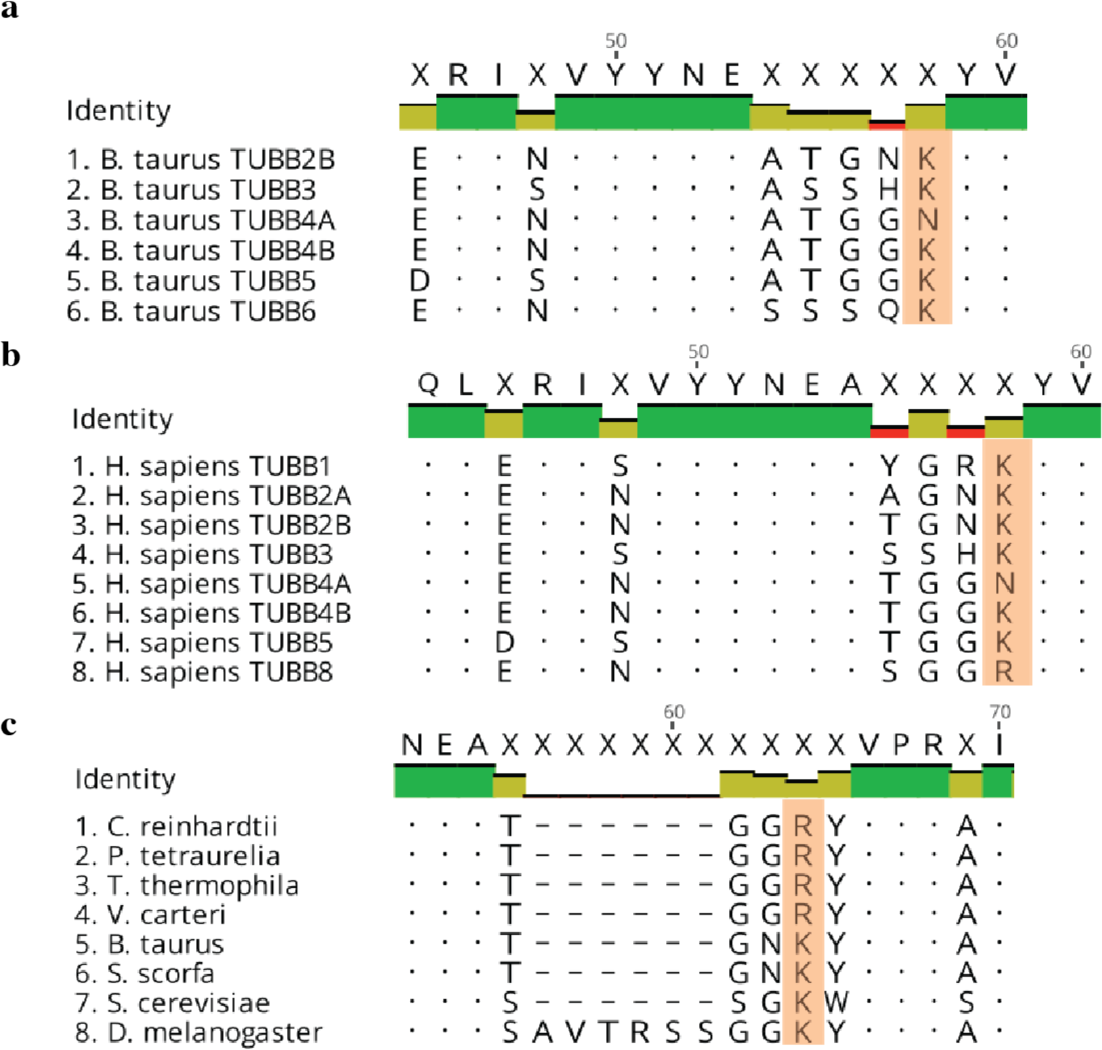
Tubulin sequence alignment for Lys-60. (**a**) Bovine tubulin isotypes include mainly lysine (K) residue at position 60 of b-tubulin, except TUBB4A that includes Asp (N) (**b**) Human tubulin isotypes include mainly lysine residue at position 60 of b-tubulin, except for TUBB4A that includes Asp (N), and TUBB8 that includes Arg (R). (**c**) Ciliates from different phylogenetic background include Arg-60, while other organisms include K-60.

**Supplementary Figure 6.**
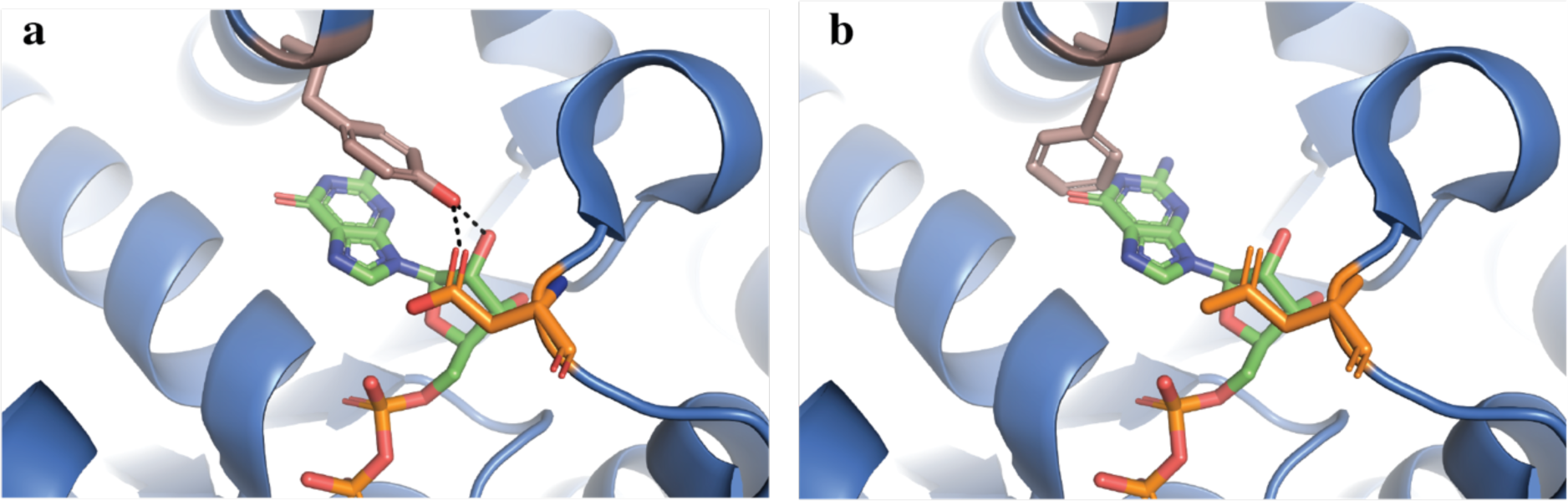
(**a**) Tyr-224 (salmon) in helix H7 at the E-site interacts with the nucleobase and Asp-179 (orange). (**b**) Substitution of Tyr-224 to Phe-224 (salmon) might change the orientation of GTP in the E-site, and consequently to affect the hydrolysis process of GTP to GDP (green). Due to the missing interactions of Phe-224, Asp-179 is more available to longitudinal interactions. Substitution was plotted on the solved protein structure for the GDP-sT2R (PDB: 3RYI) using PyMOL software (http://www.pymol.org/).

**Supplementary Figure 7.**
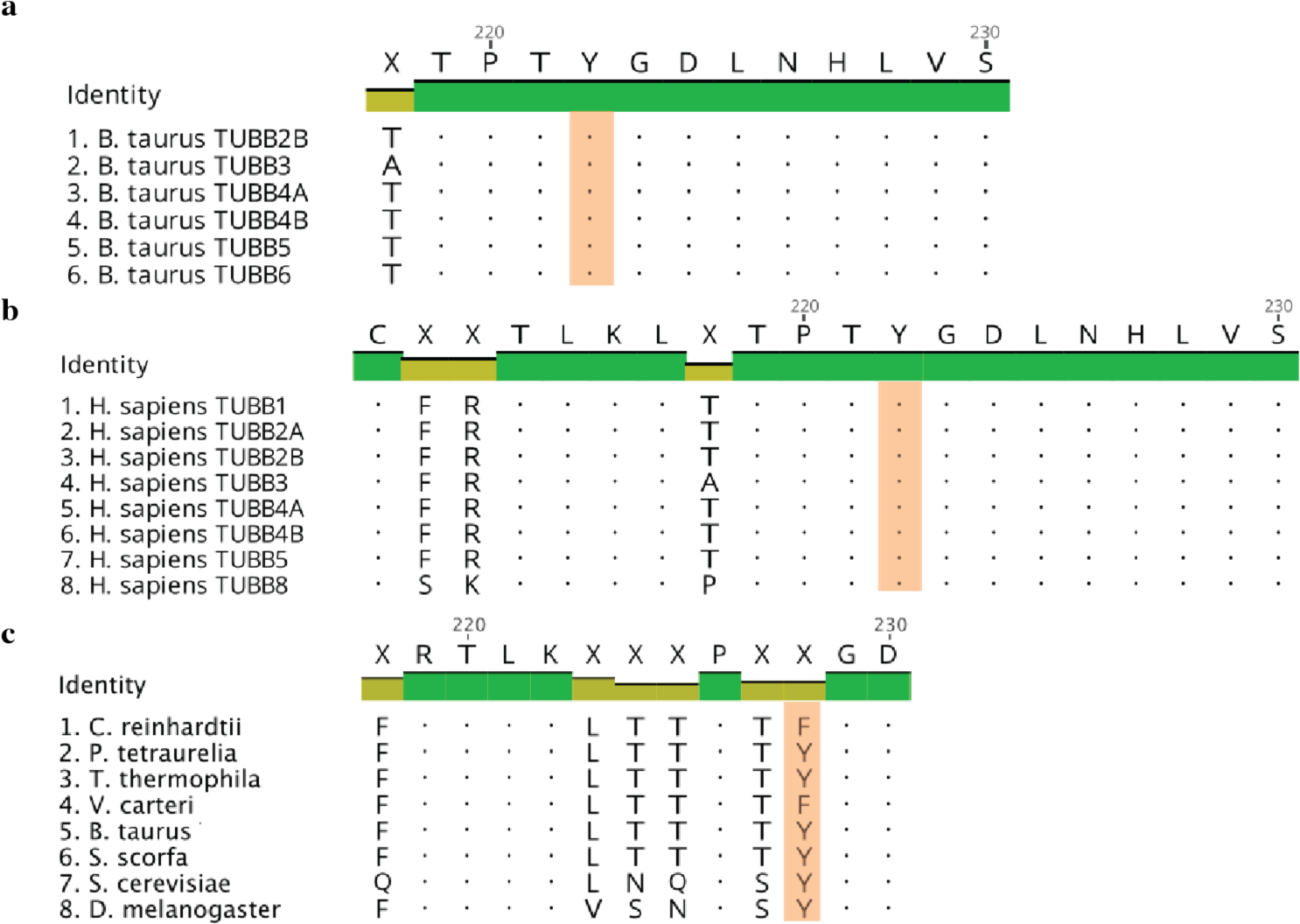
Tubulin sequence alignment for Tyr224. Bovine (**a**) and human (**b**) tubulin isotypes include Tyr (Y) residue at position 224 of b-tubulin. (**c**) Only *Chlamydomonas reinhartii* and *Volvox carteri* include Phe (F) residue at position 224, while other studied organism include Tyr-224.

**Supplementary Figure 8.**
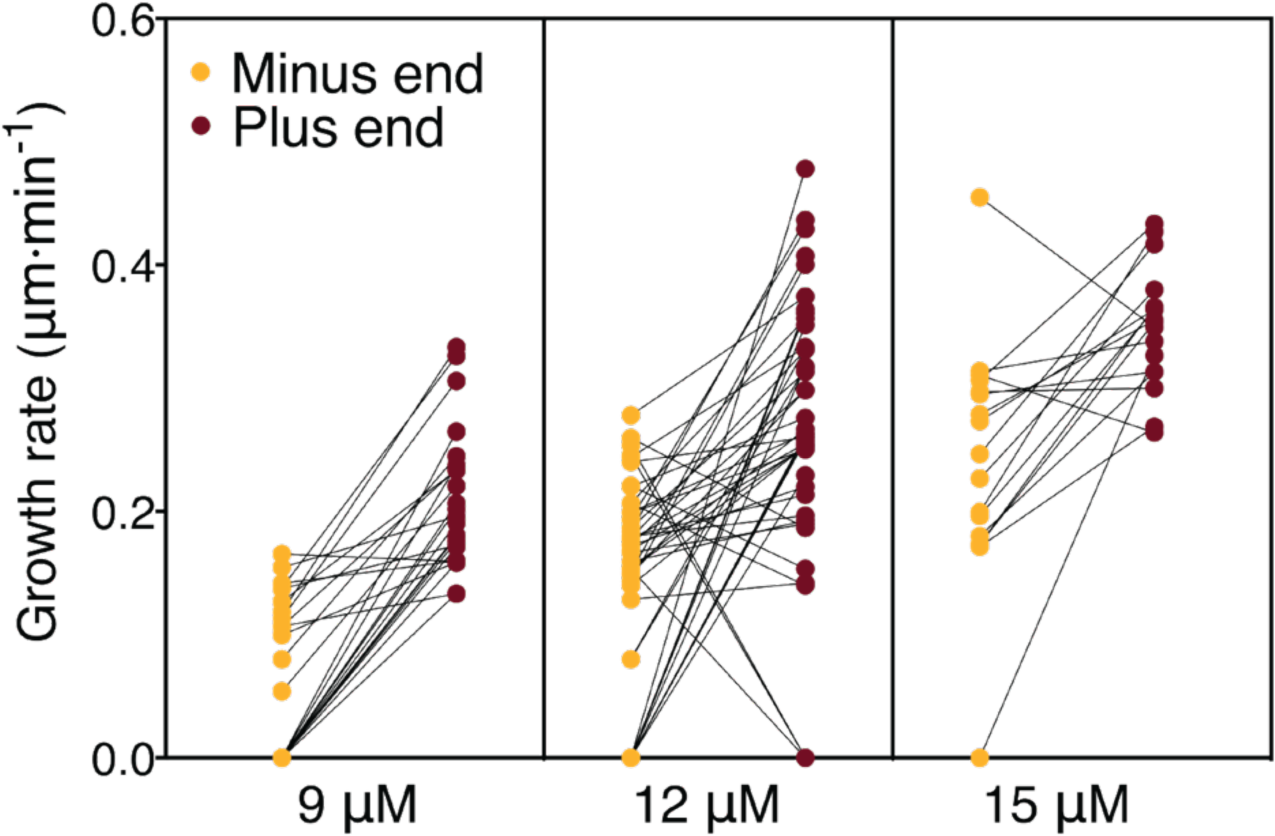
Dual-labeled GMPCPP stabilized bovine MTs seeds were used to follow the growth rate at minus end (blue) and the plus end (red) of axonemal tubulin at different tubulin concentrations.

## References

1 Carvalho-Santos, Z., Azimzadeh, J., Pereira-Leal, J. B. & Bettencourt-Dias, M. Evolution: Tracing the origins of centrioles, cilia, and flagella. J. Cell Biol. 194, 165–175 (2011).

2 Goetz, S. C. The primary cilium: a signalling centre during vertebrate development. Nat. Rev. Genet. 11, 331–344 (2010).

3 Singla, V. & Reiter, J. F. The primary cilium as the cell’s antenna: signaling at a sensory organelle. Science 313, 629–633 (2006).

4 Nicastro, D. et al. The molecular architecture of axonemes revealed by cryoelectron tomography. Science 313, 944–948 (2006).

5 Ishikawa, H. & Marshall, W. F. Ciliogenesis: building the cell’s antenna. Nat. Rev. Mol. Cell Biol. 12, 222–234 (2011).

6 Ichikawa, M. et al. Subnanometre-resolution structure of the doublet microtubule reveals new classes of microtubule-associated proteins. Nature Commun. 8, 15035 (2017).

7 Behnke, O. & Forer, A. Evidence for four classes of microtubules in individual cells. J. Cell Sci. 2, 169–192 (1967).

8 Witman, G. B., Carlson, K. & Rosenbaum, J. L. Chlamydomonas flagella. II. The distribution of tubulins 1 and 2 in the outer doublet microtubules. J. Cell Biol. 54, 540–555 (1972).

9 Mitchison, T. & Kirschner, M. Dynamic instability of microtubule growth. Nature 312, 237–242 (1984).

10 Kirschner, M. & Mitchison, T. Beyond Self-Assembly: From Microtubules to Morphogenesis. Cell 45, 329–342 (1986).

11 Desai, A. & Mitchison, T. J. Microtubule polymerization dynamics. Annu. Rev. Cell Dev. Biol. 13, 83–117 (1997).

12 Howard, J. & Hyman, A. A. Dynamics and mechanics of the microtubule plus end. Nature 422, 753–758 (2003).

13 Marshall, W. F. & Rosenbaum, J. L. Intraflagellar transport balances continuous turnover of outer doublet microtubules: implications for flagellar length control. J. Cell Biol. 155, 405–414 (2001).

14 Gadadhar, S. et al. Tubulin glycylation controls primary cilia length. J. Cell Biol. 216, 2701–2713 (2017).

15 Wloga, D. et al. TTLL3 Is a Tubulin Glycine Ligase that Regulates the Assembly of Cilia. Dev. Cell 16, 867–876 (2009).

16 Pathak, N., Austin, C. A. & Drummond, I. A. Tubulin tyrosine ligase-like genes ttll3 and ttll6 maintain zebrafish cilia structure and motility. J. Biol. Chem. 286, 11685–11695 (2011).

17 Bosch Grau, M. et al. Tubulin glycylases and glutamylases have distinct functions in stabilization and motility of ependymal cilia. J. Cell Biol. 202, 441–451 (2013).

18 Kubo, T., Hirono, M., Aikawa, T., Kamiya, R. & Witman, G. B. Reduced tubulin polyglutamylation suppresses flagellar shortness in Chlamydomonas. Mol. Biol. Cell 26, 2810–2822 (2015).

19 Kubo, T., Yanagisawa, H.-A., Yagi, T., Hirono, M. & Kamiya, R. Tubulin polyglutamylation regulates axonemal motility by modulating activities of inner-arm dyneins. Curr. Biol. 20, 441–445 (2010).

20 Suryavanshi, S. et al. Tubulin glutamylation regulates ciliary motility by altering inner dynein arm activity. Curr. Biol. 20, 435–440 (2010).

21 Alper, J. D., Decker, F., Agana, B. & Howard, J. The motility of axonemal dynein is regulated by the tubulin code. Biophys. J. 107, 2872–2880 (2014).

22 Kathir, P. et al. Molecular map of the Chlamydomonas reinhardtii nuclear genome. Eukaryot. Cell 2, 362–379 (2003).

23 Youngblom, J., Schloss, J. A. & Silflow, C. D. The two beta-tubulin genes of Chlamydomonas reinhardtii code for identical proteins. Mol. Cell. Biol. 4, 2686–2696 (1984).

24 James, S. W., Silflow, C. D., Stroom, P. & Lefebvre, P. A. A mutation in the alpha 1-tubulin gene of Chlamydomonas reinhardtii confers resistance to anti-microtubule herbicides. J. Cell Sci. 106, 209–218 (1993).

25 Hofmeister, F. Zur Lehre von der Wirkung der Salze. Arch. Exp. Pathol. Pharmacol. 24, 247–260 (1888).

26 Baldwin, R. L. How Hofmeister ion interactions affect protein stability. Biophysj 71, 2056–2063 (1996).

27 Mahamdeh, M., Simmert, S., Luchniak, A., Schäffer, E. & Howard, J. Label-free high-speed wide-field imaging of single microtubules using interference reflection microscopy. J. Microsc. 1991, 95 (2018).

28 Hill, T. L. & Carlier, M. F. Steady-state theory of the interference of GTP hydrolysis in the mechanism of microtubule assembly. Proc. Natl. Acad. Sci. USA 80, 7234–7238 (1983).

29 Gardner, M. K. et al. Rapid microtubule self-assembly kinetics. Cell 146, 582–592 (2011).

30 Walker, R. A. et al. Dynamic instability of individual microtubules analyzed by video light microscopy: rate constants and transition frequencies. J. Cell Biol. 107, 1437–1448 (1988).

31 Toso, R. J., Jordan, M. A., Farrell, K. W., Matsumoto, B. & Wilson, L. Kinetic stabilization of microtubule dynamic instability in vitro by vinblastine. Biochemistry 32, 1285–1293 (1993).

32 Lechtreck, K. F. & Geimer, S. Distribution of polyglutamylated tubulin in the flagellar apparatus of green flagellates. Cell Motil. Cytoskeleton 47, 219–235 (2000).

33 Johnson, K. A. The axonemal microtubules of the Chlamydomonas flagellum differ in tubulin isoform content. J. Cell Sci. 111, 313–320 (1998).

34 Robison, P. et al. Detyrosinated microtubules buckle and bear load in contracting cardiomyocytes. Science 352, aaf0659–aaf0659 (2016).

35 Kubo, T. & Oda, T. Electrostatic interaction between polyglutamylated tubulin and the nexin-dynein regulatory complex regulates flagellar motility. Mol. Biol. Cell 28, 2260–2266 (2017).

36 Kubo, T., Yagi, T. & Kamiya, R. Tubulin polyglutamylation regulates flagellar motility by controlling a specific inner-arm dynein that interacts with the dynein regulatory complex. Cytoskeleton (Hoboken) 69, 1059–1068 (2012).

37 McIntosh, J. R. et al. Microtubules grow by the addition of bent guanosine triphosphate tubulin to the tips of curved protofilaments. J. Cell Biol. 265, jcb.201802138 (2018).

38 Guesdon, A. et al. EB1 interacts with outwardly curved and straight regions of the microtubule lattice. Nature Cell Biol. 18, 1102–1108 (2016).

39 Chrétien, D., Fuller, S. D. & Karsenti, E. Structure of growing microtubule ends: two-dimensional sheets close into tubes at variable rates. J. Cell Biol. 129, 1311–1328 (1995).

40 Kerssemakers, J. W. J. et al. Assembly dynamics of microtubules at molecular resolution. Nature 442, 709–712 (2006).

41 Schek, H. T., Gardner, M. K., Cheng, J., Odde, D. J. & Hunt, A. J. Microtubule assembly dynamics at the nanoscale. Curr. Biol. 17, 1445–1455 (2007).

42 Zhang, R., Alushin, G. M., Brown, A. & Nogales, E. Mechanistic Origin of Microtubule Dynamic Instability and Its Modulation by EB Proteins. Cell 162, 849–859 (2015).

43 Manka, S. W. & Moores, C. A. The role of tubulin-tubulin lattice contacts in the mechanism of microtubule dynamic instability. Nat. Struct. Mol. Biol. 114, 977 (2018).

44 Nawrotek, A., Knossow, M. & Gigant, B. The determinants that govern microtubule assembly from the atomic structure of GTP-tubulin. J. Mol. Biol. 412, 35–42 (2011).

45 Tyler, K. M., Wagner, G. K., Wu, Q. & Huber, K. T. Functional significance may underlie the taxonomic utility of single amino acid substitutions in conserved proteins. J. Mol. Evol. 70, 395–402 (2010).

46 Mandelkow, E. M. Microtubule dynamics and microtubule caps: a time-resolved cryo- electron microscopy study. J. Cell Biol. 114, 977–991 (1991).

47 Nogales, E. & Wang, H.-W. Structural mechanisms underlying nucleotide-dependent self-assembly of tubulin and its relatives. Curr. Opin. Struct. Biol. 16, 221–229 (2006).

48 Brouhard, G. J. & Rice, L. M. The contribution of αβ-tubulin curvature to microtubule dynamics. J. Cell Biol. 207, 323–334 (2014).

49 Ayaz, P., Ye, X., Huddleston, P., Brautigam, C. A. & Rice, L. M. A TOG:αβ-tubulin complex structure reveals conformation-based mechanisms for a microtubule polymerase. Science 337, 857–860 (2012).

50 Pecqueur, L. et al. A designed ankyrin repeat protein selected to bind to tubulin caps the microtubule plus end. Proc. Natl. Acad. Sci. USA 109, 12011–12016 (2012).

51 Rice, L. M., Montabana, E. A. & Agard, D. A. The lattice as allosteric effector: Structural studies of alpha beta- and gamma-tubulin clarify the role of GTP in microtubule assembly. Proc. Natl. Acad. Sci. USA 105, 5378–5383 (2008).

52 Vitre, B. et al. EB1 regulates microtubule dynamics and tubulin sheet closure in vitro. Nature Cell Biol. 10, 415–421 (2008).

53 Müller-Reichert, T., Chrétien, D., Severin, F. & Hyman, A. A. Structural changes at microtubule ends accompanying GTP hydrolysis: information from a slowly hydrolyzable analogue of GTP, guanylyl (alpha, beta)methylenediphosphonate. Proc. Natl. Acad. Sci. USA 95, 3661–3666 (1998).

54 Honnappa, S., Cutting, B., Jahnke, W., Seelig, J. & Steinmetz, M. O. Thermodynamics of the Op18/Stathmin-Tubulin Interaction. J. Biol. Chem. 278, 38926–38934 (2003).

55 Barbier, P. et al. Stathmin and interfacial microtubule inhibitors recognize a naturally curved conformation of tubulin dimers. J. Biol. Chem. 285, 31672–31681 (2010).

## References

56 Alper, J., Geyer, V., Mukundan, V. & Howard, J. Reconstitution of flagellar sliding. Meth. Enzymol. 524, 343–369 (2013).

57 Gorman, D. S. & Levine, R. P. Cytochrome f and plastocyanin: their sequence in the photosynthetic electron transport chain of Chlamydomonas reinhardi. Proc. Natl. Acad. Sci. USA 54, 1665–1669 (1965).

58 Castoldi, M. & Popov, A. V. Purification of brain tubulin through two cycles of polymerization-depolymerization in a high-molarity buffer. 32, 83–88 (2003).

59 Gell, C. et al. Microtubule Dynamics Reconstituted In Vitro and Imaged by Single-Molecule Fluorescence Microscopy. Methods Cell Biol. 95, 221–245 (2010).

60 Witman, G. B. Isolation of Chlamydomonas Flagella and Flagellar Axonemes. Meth. Enzymol. 134, 280–290 (1986).

61 LeDizet, M. & Piperno, G. Detection of acetylated alpha-tubulin by specific antibodies. Meth. Enzymol. 196, 264–274 (1991).

62 Wolff, A. et al. Distribution of glutamylated alpha and beta-tubulin in mouse tissues using a specific monoclonal antibody, GT335. 59, 425–432 (1992).

63 Ruhnow, F., Zwicker, D. & Diez, S. Tracking Single Particles and Elongated Filaments with Nanometer Precision. Biophys. J. 100, 2820–2828 (2011).

64 Tarantino, N. et al. TNF and IL-1 exhibit distinct ubiquitin requirements for inducing NEMO-IKK supramolecular structures. J. Cell Biol. 204, 231–245 (2014).

